# N-myristoyltransferase Inhibitors as Candidate Broad-Spectrum Antivirals to Treat Viral Infections Promoted by Immunosuppression Associated with JAK Inhibitors Therapy

**DOI:** 10.1101/2025.06.20.660686

**Authors:** Haydar Witwit, Juan Carlos de la Torre

## Abstract

The use of Janus kinase (JAK) inhibitors in the clinic has been expanded significantly during the last decade. However, the immunosuppressive effects of JAK inhibitors, via modulation of key innate cellular signaling pathways, can predispose treated patients to infections, and can also result in reduced control of silent infections and increased risk of reactivation of opportunistic infections. Thus, the JAK inhibitor ruxolitinib, approved for the treatment of myelofibrosis and polycythemia vera, has been shown to exerts a proviral activity during infection with different viruses. Therefore, the clinical relevance of developing antiviral treatments that can be effective in the presence of JAK inhibitors. N-terminal myristoyl transferase (NMT) inhibitors have been shown to exhibit potent antiviral activity against different viruses. Here we document that in the presence of ruxolitinib, NMT inhibitors retain their potent antiviral activity against different viruses, including HSV-1. Our findings support that NMT inhibitors should be explored as therapeutics to treat viral infections associated with immunosuppression caused by treatments with JAK inhibitors.

## Introduction

Ruxolitinib (Rux), the first approved JAK inhibitor, in 2011, was initially indicated to treat intermediate-and high-risk myelofibrosis cases in primary, post-polycythemia, and post-essential thrombocythemia patients. Subsequently, the approval of Rux has been expanded to treat polycythemia vera (2014) [1], which was followed by its approval for use in acute graft-versus-host disease (2019) [2], then chronic graft-versus-host disease (2020) [3], and recently it is undergoing a clinical trial to treat multiple myeloma[4]. Nine FDA-approved JAK inhibitors (JAKi) are currently in use in clinical setting with broad covering indications including rheumatoid arthritis (RA) (tofacitinib, baricitinib, and upadacitinib), ulcerative colitis (UC) (tofacitinib, and upadacitinib), atopic dermatitis (abrocitinib, upadacitinib, and opzelura), psoriatic arthritis (PsA) (tofacitinib, and upadacitinib), and additional JAKi have entered clinical trials.

The expected future impact of JAKi in the clinic and health care system is illustrated by the estimated over fifty million patients suffering from RA in USA[5] with an observed high percentage of non-compliance to medications in the pre-JAKi era due to limited efficacy and significant side effects[6], and current guidelines favoring JAKi-based combination therapy to start in phase II, when phase I therapies with methotrexate, leflunomide or sulfasalazine in combination with corticosteroids fails to show clinical improvement. Likewise, JAKi are also established as an RA phase III therapy[7]. However, the use of JAKi has the common side effect of attenuating the type I interferon (T1IFN) response, a main player of the cell innate immune response to infection, which can result in immunosuppression and facilitate de novo infections, or reactivation, of viruses that are normally controlled by the T1IFN response such as varicella zoster [8, 9] [10]. These situations require pausing JAKi therapy until infection is controlled by antiviral therapy [11], which can be jeopardized by the emergence of antiviral resistant viral strains [11, 12] that can be also able to counteract the antiviral activity of the T1IFN response [8, 13–16]. Preventive vaccination can reduce varicella zoster incidence by ∼50% in patients treated with JAKi, but the other ∼50% of cases remain liable to the immunosuppressive effects of JAKi, which increase susceptibility to viral infections. Hence, the clinical significance of developing antiviral treatments that can be effective in the presence of JAKi. N-myristoyltransferases (NMTs) catalyze the covalent attachment of myristic acid to the N-terminal glycine of substrate proteins—a modification crucial for membrane localization and function of many viral and host proteins [17–21]. Initially studied in the context of parasitic diseases [22–24] and cancer [25], NMT inhibitors have more recently emerged as indirect antiviral due to their role in interrupting replication and assembly of diverse viruses [18, 21, 26, 27].

Here, we present evidence that NMT inhibitors retain their potent antiviral activity in the presence of Rux. Moreover, by targeting a host cell protein, NMT inhibitors pose a higher genetic barrier to the emergence of drug resistant viral variants [18, 28].

## Materials and methods

### Cells and viruses

Homo sapiens A549 (ATCC CCL-185) and Vero E6 (ATCC CRL-1586) (*Chlorocebus aethiops*) cell lines were maintained in Dulbecco’s modified Eagle medium (DMEM) (ThermoFisher Scientific, Waltham, MA, USA) containing 10% heat-inactivated fetal bovine serum (FBS), 2 mM L-glutamine, 100 μg/mL streptomycin, and 100 U/mL penicillin. Wild type (WT) lymphocytic choriomeningitis virus (LCMV) and its NP(D382A) mutant form[29], a recombinant vaccinia virus expressing the nano luciferase and GFP reporter genes (rVACV-Nluc/GFP)[30], have been described.

### Compounds and antibodies

Ruxolitinib (Cat. No.: S1378) was purchased from Selleck Chem. DDD85646 (Cat. No.: HY-103056) was purchased from MedChem Exp (NJ, USA). Rat monoclonal antibody VL4 against LCMV nucleoprotein (NP) was purchased from Bio C Cell (West Lebanon, NH, USA). Anti-rat Alexa Fluor 488 and 568 were conjugated with VL4 to recognize NP.

### Statistical analyses

Ordinary two-way ANOVA statistical test with recommended Tukey correction for multiple comparison tests were performed to detect statistical significance for differences in virus titer, the multiplicity average P value for each comparison, in each test, was reported with alpha threshold and confidence interval established at 0.05 and 95% respectively. Statistically significant probability values were represented as follows: ns *p* > 0.05, ** *p* < 0.01, *** *p* < 0.001, **** *p* < 0.0001. The analyses were performed GraphPad Prism (GraphPad Software, Boston, MA USA), v10 (Prism10).

## Results

### Effect of the JAKi Rux on multiplication of LCMV

To assess the effect of Rux on multiplication of viruses with different degree of susceptibility to the antiviral activity of T1IFN, we examined the effect of Rux on multiplication of the prototypic mammarenavirus lymphocytic choriomeningitis virus (LCMV) that poses a lethal threat to immunocompromised transplantation patients[31, 32]. We compared the effect of Rux on wild type (WT) LCMV, which is able to counteract the host cell T1IFN response, and a mutant form of LCMV (rLCMV /NP(D382A), where mutation D382A in the virus nucleoprotein (NP) interferes with LCMV’s ability to counteract the T1IFN response[16, 33]. We treated A549 cells with Rux starting at 24 hours (h) pre-infection or 90 minutes post-infection with either WT or NP(D382A) LCMV, and at 72 h post-infection (pi) we fixed the cells and determined the number of infected cells by IF using a NP-specific antibody (Fig 1). Consistent with published findings WT, but not NP(D382A), LCMV exhibited robust multiplication in A549 cells. Treatment with Rux did not significantly affect multiplication of LCMV WT, but alleviated the restricted multiplication of NP(D382A) in A549 cells.

**Figure 1.**
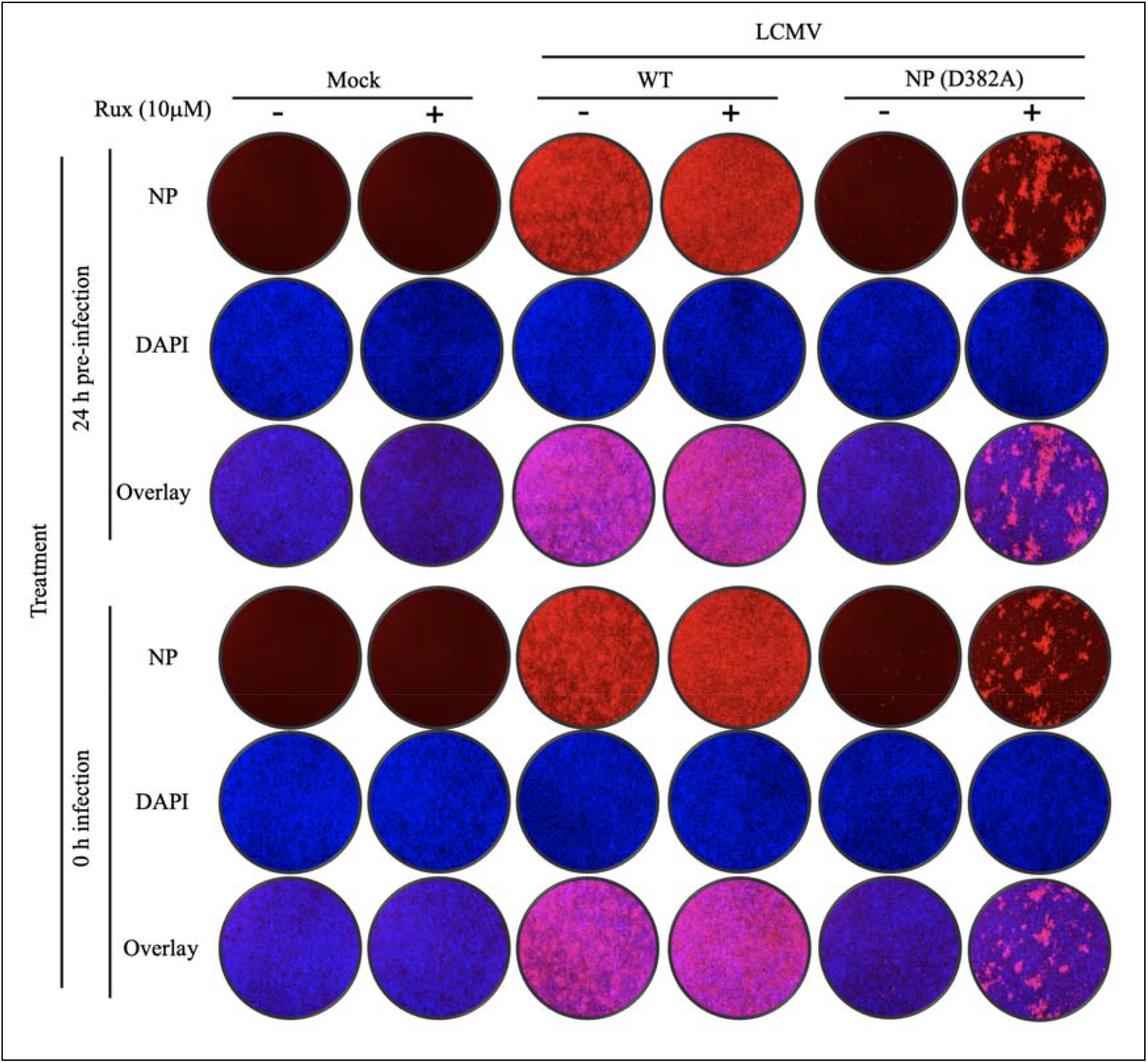
The JAKi Rux promotes multiplication of rLCMV/NP(D382A) in A549 cells. A549 cells were seeded at 2.0 × 10^5^ cells/well into a 24-well plate, infected with rLCMV WT (MOI 0.01) or NP(D382A) mutant (MOI 0.05) and treated with Rux (10 μM) alone or in combination with NMTi (5 µM). Cells were fixed at the indicated h pi and infected cells identified by staining with the rat monoclonal antibody to NP VL4. Cell nuclei were visualized by DAPI staining. Immunofluorescence images (4X magnification) were taken using Keyence BZ-X710.

### Activity of NMT inhibitors in the presence of the JAKi Rux

NMT inhibitors (NMTi) have been shown to potently inhibit mammarenaviruses[18, 34]. We investigated whether the NMTi DDD85646 retained its potent antiviral activity against LCMV in cells treated with Rux. For this, we examined the effect of NMTi on the multi-step growth kinetic of WT and NP(D382A) LCMV in the absence or presence of Rux (Fig. 2). As predicted, Rux-mediated inhibition of the T1IFN pathway resulted in readily detected multiplication of rLCMV/NP(D382A) in A549 cells, which was inhibited in the presence of NMTi (Fig. 2). These findings support an T1IFN-independent anti-mammarenaviral activity of NMTi. In cells infected with rLCMV/NP(D382A) and treated with Rux + NMTi, we detected similarly low numbers of NP^+^ cells at 24, 48 and 72 h pi by IF, whereas titers of infectious progeny in cell culture supernatants (CCS) were below detection at 48 and 72 h pi. These findings are consistent with our published results showing that NMTi do not significantly affect LCMV replication and transcription, but rather target the roles of the LCMV Z matrix protein in viral assembly and budding processes[18], together with the reported long half-life (∼ 72 h) of NP[35]. The observed decrease in the intensity of NP staining over time in rLCMV/NP(D382A) infected cells in the absence of JAKi treatment likely reflects the NP decay process magnified by the T1IFN mediated blockade of virus multiplication. We also confirmed that cell density, which could influence cytokine production [36, 37], did not influence infection in presence of Rux, or the antiviral activity of NMTi.

**Figure 2.**
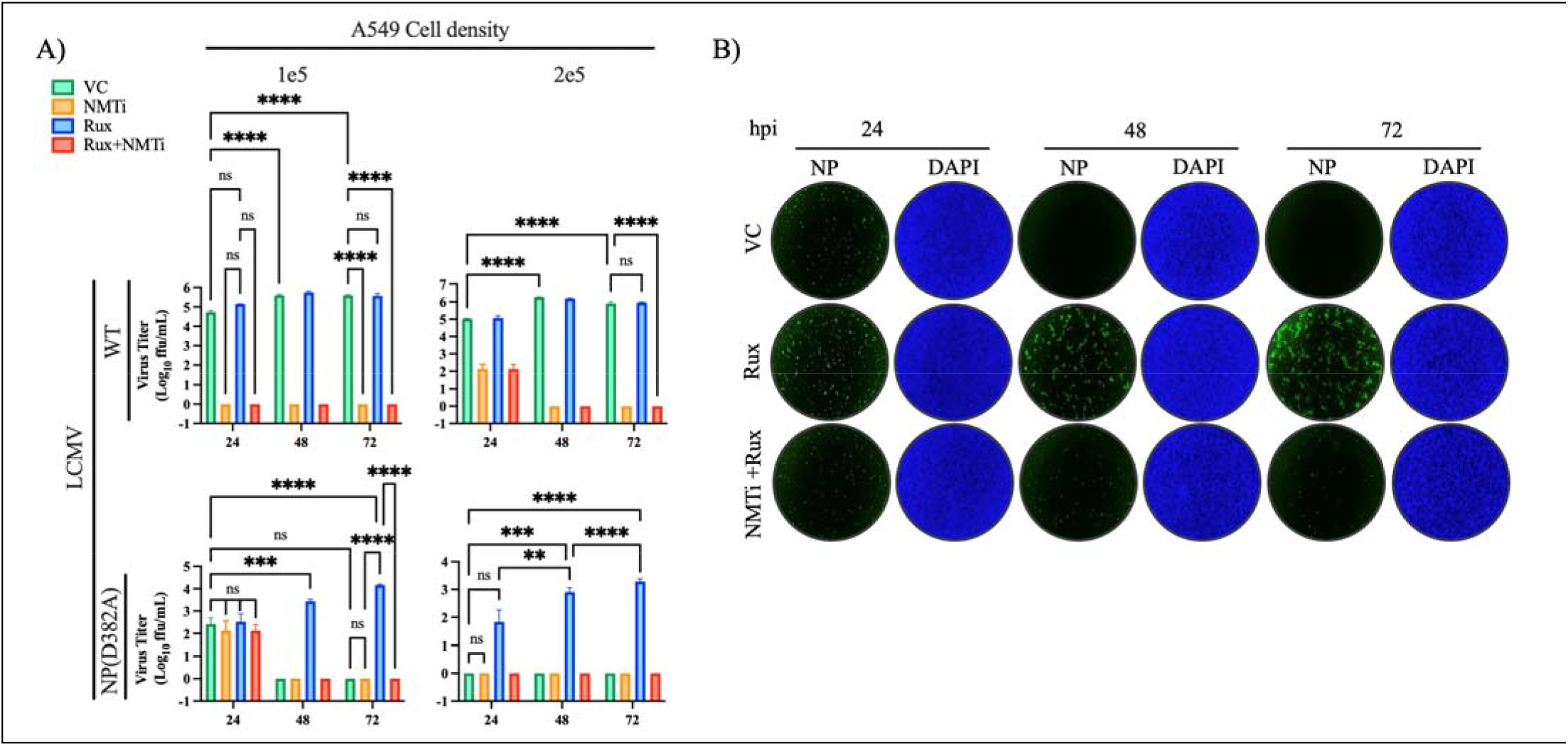
Effect of NMTi on multiplication of LCMV in A549 cells in the presence of the JAKi Rux. A549 cells were seeded at 1.0 (1e5) or 2.0 × 10^5^ (2e5) cells/well into a 24-well plate, infected with rLCMV WT or NP(D382A) mutant at MOI 0.01 and 0.05, respectively, and treated with Rux (10 μM) alone or in combination with NMTi (5 µM). Cell culture supernatants (CCS) were collected at the indicated time points and titers of infectious virus determined by FFA (**A)**. Cells infected with rLCMV/NP(D382A) were fixed at the indicated h pi and infected cells identified by staining with the rat monoclonal antibody VL4 to NP VL4 (**B**). Nuclei were identified by DAPI staining. Immunofluorescence images (4X magnification) were taken using Keyence BZ-X710.

### Effect of NMTi on Vaccinia virus (VACV) multiplication in the presence of the JAKi Rux

VACV can be used as a valid surrogate for the assessment of antivirals effective against monkeypox virus (MPXV)[28]. We have shown that NMTi potently inhibit multiplication of VACV[27, 28]. We examined whether the potent anti-VACV activity of NMTi was retained in the presence of Rux (Fig 3). NMTi was able to inhibit multiplication of VACV in presence of Rux as determined by the numbers of infected cells detected by IF (Fig. 3A) and production of VACV infectious progeny (Fig. 3B) in a multi-step growth kinetics assay.

**Figure 3.**
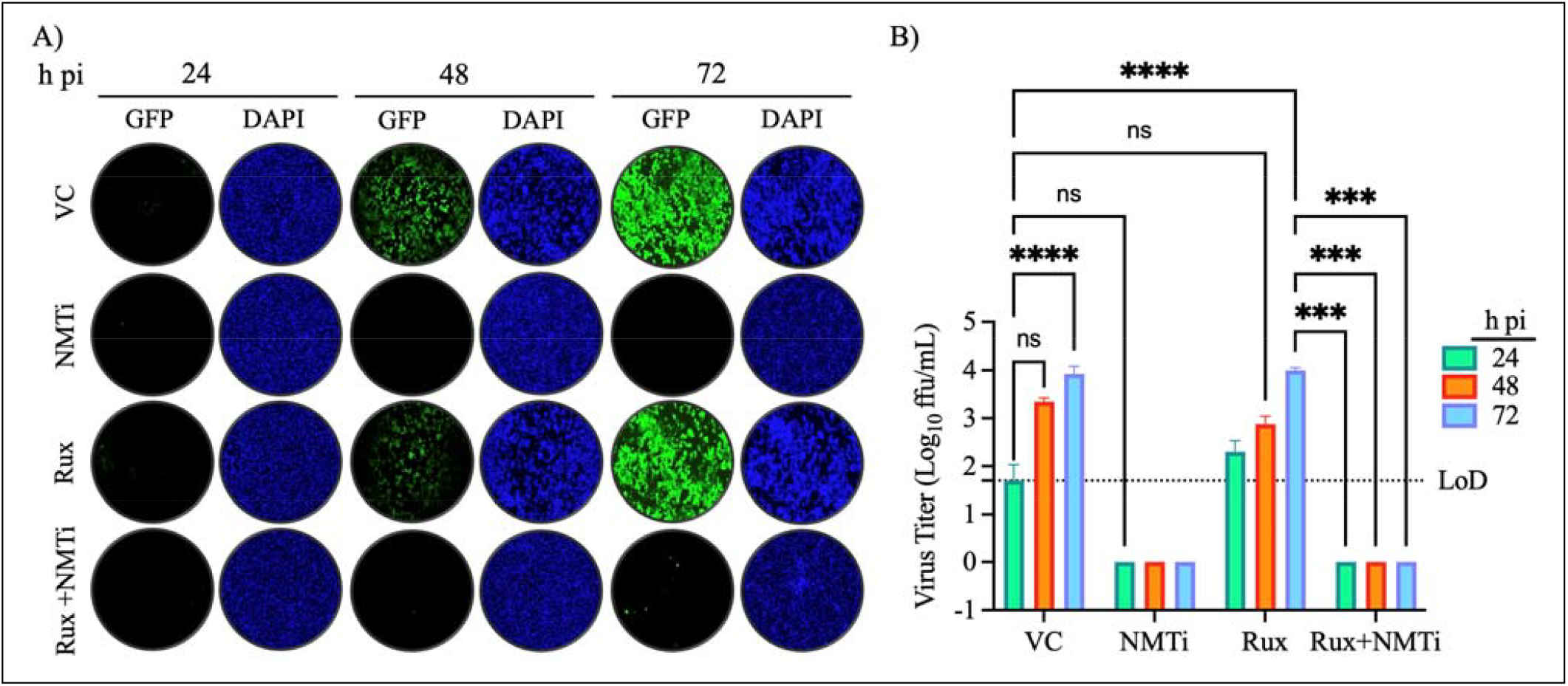
Effect of NMTi on multiplication of rVACV-Nluc/GFP in A549 cells in the presence of Rux. A549 cells were seeded at 2.0 × 10^5^ cells/well into a 24-well plate, infected with rVACV-Nluc/GFP (MOI 0.01) and treated with Rux (10 μM) alone or in combination with NMTi (5 µM). Cells were fixed at the indicated h pi and stained with DAPI. Infected cells were identified by the expression of GFP (**A**). Cell culture supernatants (CCS) were collected at the indicated time points and titers of infectious virus determined by FFA (**B**). Immunofluorescence images (4X magnification) were taken using Keyence BZ-X710.

### Effect of NMTi on HSV-1 multiplication in the presence of the JAKi Rux

Evidence indicates that several HSV-1 proteins undergo post-translational fatty acylation [38– 41], including myrisotylation[39–41], but the effect of NMTi on HSV-1 multiplication has not been investigated. We found that HSV-1 multiplication was effectively inhibited by treatment with NMTi (Fig. 4). Rux treatment has been shown to increase the risk of HSV-1 and varicella zoster (VZV) recurrence[42–45]. We therefore examined whether the anti-HSV-1 activity of NMTi was affected in the presence of Rux. For this, we treated A549 cells with Rux (10 µM) for 24 hours, followed by infection with HSV-1 in the presence of Rux and presence or absence of NMTi. We found that HSV-1 multiplication was effectively inhibited by NMTi in the presence of Rux in a multi-step growth kinetics assay (Fig. 4).

**Figure 4.**
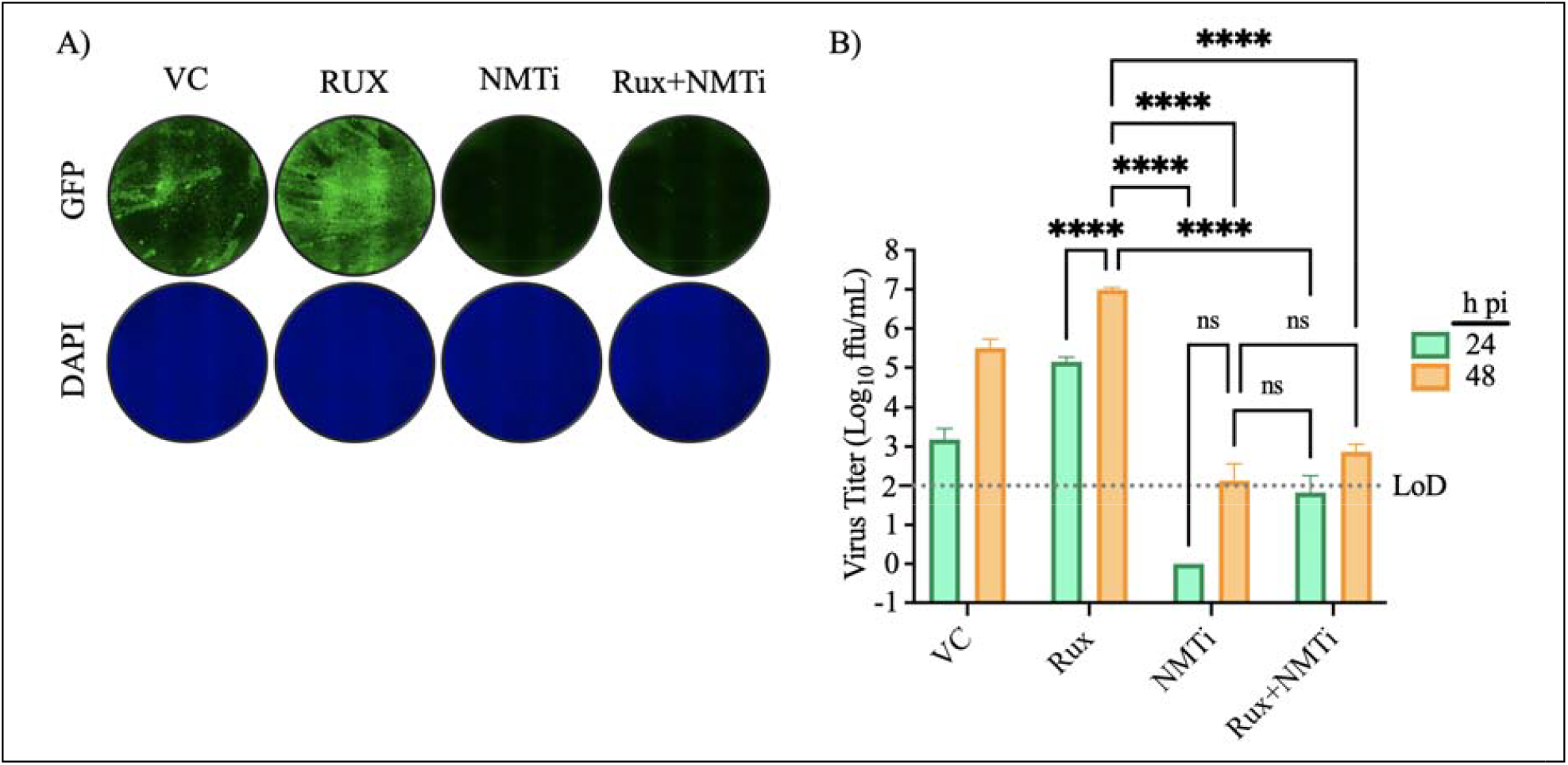
Effect of NMTi on multiplication of the HSV-1 in A549 cells in the presence of Rux. A549 cells were seeded at 2.5 × 10^5^ cells/well into a 24-well plate, infected with rHSV-1-GFP (MOI 0.005) and treated with Rux (10 μM) alone or in combination with NMTi (5 µM). Cells were fixed at 48 h pi and stained with DAPI. Infected cells were identified by the expression of GFP (**A**). Cell culture supernatants (CCS) were collected at the indicated time points and titers of infectious virus determined by FFA (**B**). Immunofluorescence images (4X magnification) were taken using Keyence BZ-X710.

## Discussion

Increased susceptibility to viral infection is a common side effect of JAKi[46, 47], which represents a challenge to the clinical management of the affected patients, as it requires pausing the JAKi therapy until the viral infection is controlled by antiviral therapy[48], with the risk of the emergence and selection of drug resistant viral variants[49–53] associated with morbidity and mortality that can reach case fatality rates up to 28%[54–56] [57, 58]. The development of host-targeted antivirals that are active in the presence of JAKi can facilitate the implementation of clinical interventions to effectively control these viral infections without interrupting the JAKi therapy and minimizing the risk of selection of drug-resistant viruses.

Consistent with published findings[42, 43, 48], the JAKi Rux exhibited a proviral activity during infection with different viruses including LCMV, known to be a threat to immunocompromised patients, VACV, a surrogate of MPXV, and HSV-1, a common human latent infection that is also used as a surrogate of varicella zoster virus. Rux treatment enhanced multiplication of rLCMV/NP(D382A), which is normally controlled by the host cell innate immunity response to infection. NMT inhibitors have been shown to potently inhibit multiplication of different viruses including mammarenaviruses [18, 34], and VACV[18, 26–28]. Early studies indicated that myristoylated HSV-1 protein UL11 was not essential for the viral multiplication [59]. However, a recent study showed that the myristoylation of UL11 can increase HSV-1 pathogenicity, and proposed myristoylation as a druggable target to counteract HSV-1 infections [60]. Consistent with these findings, our results demonstrated that NMTi reduced production of infectious HSV-1 progeny by more than four logs at 48 h pi, and that this potent antiviral activity was not affected in the presence of the JAKi Rux.

NMTi have been historically associated with low specificity and significant toxicity, thus limiting the interest of exploring their potential as antivirals. However, the generation of novel NMTi exhibiting high specificity and low associated toxicity has reignited their interest as potential antivirals [23, 24, 26, 61]. Notably, a recent human phase I trial has shown that the oral, highly bioavailable small-molecule NMTi PCLX-001 is safe and well tolerated at concentrations that result in PCLX-001 plasma concentrations over its EC_90_ [62]. Progress on NMTi-based cancer therapies can facilitate the repurposing of NMTi as antiviral agents to treat some of the infections associated with JAKi-based therapies.

## Author contributions

Conceptualization, H.W. and J.C.d.l.T.; Formal analysis, H.W. and J.C.d.l.T.; Investigation, H.W.; Methodology, H.W.; Project administration, H.W.; Resources, J.C.d.l.T.; Software, H.W.; Validation, H.W.; Visualization, H.W.; Writing—original draft, H.W. and J.C.d.l.T.; Writing—review and editing, H.W. and J.C.d.l.T. All authors have read and agreed to the published version of the manuscript.

## Ethical approval

Not applicable.

## Consent for publication

Not applicable.

